# Stretchable mesh microelectronics for the biointegration and stimulation of neural organoids

**DOI:** 10.1101/2022.06.10.495715

**Authors:** Thomas L. Li, Yuxin Liu, Csaba Forro, Levent Beker, Zhenan Bao, Bianxiao Cui, Sergiu P. Paşca

**Author notes:** These authors contributed equally (Thomas L. Li, Yuxin Liu).

## Abstract

Advances in tridimensional (3D) culture approaches have led to the generation of organoids that recapitulate cellular and physiological features of domains of the human nervous system. Although microelectrodes have been developed for long-term electrophysiological interfaces with neural tissue, studies of long-term interfaces between microelectrodes and free-floating organoids remain limited. In this study, we report a stretchable, soft mesh electrode system that establishes an intimate *in vitro* electrical interface with human neurons in 3D organoids. Our mesh is constructed with poly(3,4-ethylenedioxythiophene) polystyrene sulfonate (PEDOT:PSS) based electrically conductive hydrogel electrode arrays and an elastomeric poly(styrene-ethylene-butadiene-styrene) (SEBS) as the substrate and encapsulation materials. This mesh electrode can maintain stable electrochemical impedance in buffer solution under 50% compressive and 50% tensile strain. We have successfully cultured pluripotent stem cell-derived human cortical organoids (hCO) on this polymeric mesh for more than 3 months and demonstrated that organoids readily integrate with the mesh. Using simultaneous stimulation and calcium imaging, we show that electrical stimulation through the mesh can elicit intensity-dependent calcium signals comparable to stimulation from a bipolar stereotrode. This platform may serve as a tool for monitoring and modulating the electrical activity of *in vitro* models of neuropsychiatric diseases.

## Introduction

Mesh electrodes are an emerging platform for chronic electrophysiological interfaces with brain tissue^1,2^. Unlike traditional multi-electrode arrays or shank probes, which are made of stiff materials such as silicon, mesh electrodes are comprised of flexible conductive interconnects and electrodes encapsulated by insulating polymeric materials. Mesh electrodes have been shown to achieve a stable long-term interface for several reasons. First is their low bending stiffness: by having thin layers, they may more easily conform to neural tissue to potentially minimize foreign body interactions^3^. Second, mesh electrodes exclude far less volume than other techniques such as solid electrode inserts. Mesh electrodes can be made thinner than 1 μm and have been shown to expand and twist open after injection into liquid solution^4,5^.

One potential area of applications for mesh electrodes is to stimulate and monitor the emergence of electrical activity in 3D neural organoids. Neural organoids begin as 3D aggregates of human induced pluripotent stem cells (hiPSCs). Over time, the hiPSC-derived differentiated cells self-organize into 3D structures recapitulating some aspects of domains of the developing neural axis^6^. These organoids, or their combination to form assembloids, can be used to study early neural network development, the cell-cell interactions that give rise to cellular organization and circuit assembly in the nervous stem, and disease-related phenotypes in patient-derived cultures. As a result, they have been used to model genetic form of neurodevelopmental diseases such as Rett syndrome^7^, Timothy syndrome^8^, 22q11.2 deletion syndrome^9^, tuberous sclerosis^10^ or Miller-Dieker Syndrome^11,12^. Neural organoids have also been used to model various insults to the developing brain, including hypoxia^13^ and Zika virus infection^14^.

To date, electrophysiological measurements on brain organoids have been primarily performed using patch clamp^9,14–20^, multi-electrode arrays^20-22^, silicon probes^20,24^, or glass pipettes^25^. The majority of these recordings were performed with acute and destructive techniques, such as slicing, while others required plating the organoid onto a planar surface, making it difficult to study the same organoid over many timepoints. Chronic electrophysiology is crucial for understanding their developmental stages, characterizing functional connections, and for monitoring effects of psychiatric drugs or other interventions. Several studies have employed mesh electrodes to interface with organoids, using either a mesh based on parylene -C^26^ or SU-8^27^. While these materials are flexible at certain thicknesses, they are not intrinsically stretchable and are made with relatively high modulus materials—they are limited in their ability to adapt to an organoid that grows over time. While the impact of mesh modulus on their interactions with growing organoids is unknown, it has been well established that substrate modulus can influence cell behavior, signaling pathway and cell differentiation^28-30^. Furthermore, less damage may be expected from small vibrations and mechanical disturbance during growth. Therefore, it is of interest to develop soft and stretchable meshes.

In this study, we present an intrinsically stretchable mesh electrode system made from conducting polymer hydrogel poly(3,4-ethylenedioxythiophene) polystyrene sulfonate (PEDOT:PSS) and elastic poly(styrene-ethylene-butadiene-styrene) (SEBS) substrate and encapsulation material^31^. Unlike meshes made from polymers such as SU-8, mesh electrodes made from PEDOT:PSS and SEBS are intrinsically stretchable and maintain a high level of electrical performance under significant strain. We successfully cultured human cortical organoids (hCOs) on these mesh electrodes for more than 3 months. While electrical recording from neural organoids using these meshes requires additional work, we demonstrate that electrical stimulation delivered through the meshes can evoke calcium signals, and that varying stimulation parameters, such as current, pulse number, and frequency, can affect the calcium response. Furthermore, we demonstrate that these stimulation results are comparable to stimulations delivered using a bipolar tungsten/iridium electrode.

## Results

We designed the mesh to be stretchable through both material composition and geometric design. Both electrode and interconnect were made of stretchable conducting polymer hydrogel using poly(3,4-ethylenedioxythiophene) polystyrene sulfonate (PEDOT:PSS) physically crosslinked due to addition of an ionic liquid (IL) additive and subsequent removal of the IL, as shown in our previous work^31^. The top and bottom insulating layers were also made of a stretchable polymer, poly(styrene ethylene butylene styrene) (SEBS) (**Fig. 1a, Fig. S1**). We added a thin layer of gold above the PEDOT:PSS interconnect to further reduce the resistance of the high-aspect-ratio interconnect. The materials were patterned into a rectangular mesh with diagonal cross-connections and an overall 5 μm thickness (**Fig. 1b**). Directly spin coating photoresist on polymeric electronic materials causes swelling and degradation of electronic properties. Therefore, the PEDOT:PSS and SEBS were patterned by photolithography with a metal-based hard mask as a protective layer between the electronic materials and photoresist. The porous structure of the mesh and the top electrode openings were formed through inductively coupled plasma reactive ion etching (ICP-RIE). The mesh was fabricated on a silicon wafer with a sacrificial water-soluble dextran layer, allowing for the lift-off of the mesh in aqueous solutions (**Fig. 1c**). Each electrode consisted of an exposed PEDOT:PSS electrode with a diameter of 50 μm (**Fig. 1d, 1e**). Electrical characterization of the mesh electrode system shows that each electrode has relatively low impedance. We measured the impedance of a sample 16-electrode mesh in PBS using an Intan Stim/Recording controller. The functional electrodes in this mesh had an average impedance of 55.8 ± 19.7 kΩ (**Fig 2a**). One electrode had unusually high impedance at 1 kHz (7.6 MΩ) and two channels had low impedance (0.81 and 7.8 kΩ) which likely correspond to disconnected or damaged electrodes. Impedances were measured at frequencies ranging from 20 Hz to 5 kHz, indicating that overall impedance declines as frequency increases (**Fig. 2a**). Excluding obviously damaged electrodes (> 1 MΩ), we measured an average impedance of 20.1± 36.1 kΩ at 1 kHz across 118 electrodes in 8 meshes from 4 separate batches. The root mean square (RMS) noise of the working electrodes in PBS was measured to be 5.0 ± 0.7 μV. To investigate electrical stimulation performance, we measured current density under a bipolar pulsed voltage (25mV - 1.5V) at 50 Hz. (**Fig. 2c**). Based on these results, the mesh can deliver a maximum current of 131 μA within the water splitting voltage window (1.23 V), which corresponds to a current density of 1.67 A/cm^2^. The high current density and low impedance is a result of both the PEDOT:PSS and the nanoporous PEDOT:PSS interconnect filled with electrolyte, which gives high electrochemical surface area as demonstrated previously^31^. We performed mechanical simulations to investigate the hot spots of mechanical strain in the mesh. First, we measured the stress-strain curves of a SEBS thin film at various strain rates, showing the viscoelastic nature of the material (**Fig. S2**). These results were then used to simulate mechanical strain in the mesh design. Upon biaxial stretching of 50%, we observed that the crossings are subject to more strain than the surrounding structures (**Fig 2d**). Consequently, the crossings are also the locations of maximal stress, reaching values around 40 kPa (**Fig 2d**). For the mesh dimensions used experimentally, this value of stress equates to a weight of less than 0.2 g required to elongate the mesh by 50%, showing that it is indeed very soft and highly stretchable with minimal forces. Critically, the electrical performance of the mesh is not affected by strain. The impedance of the electrodes, measured between 20 to 5000 Hz, did not significantly change under 50% strain in either compression or stretch (**Fig 2e**, two-way ANOVA, p = 0.921 for strain). This stability and stretchability allowed the mesh to easily deform in culture, as demonstrated by manipulation with metal forceps (**Fig. 2f**).

**Figure 1:**
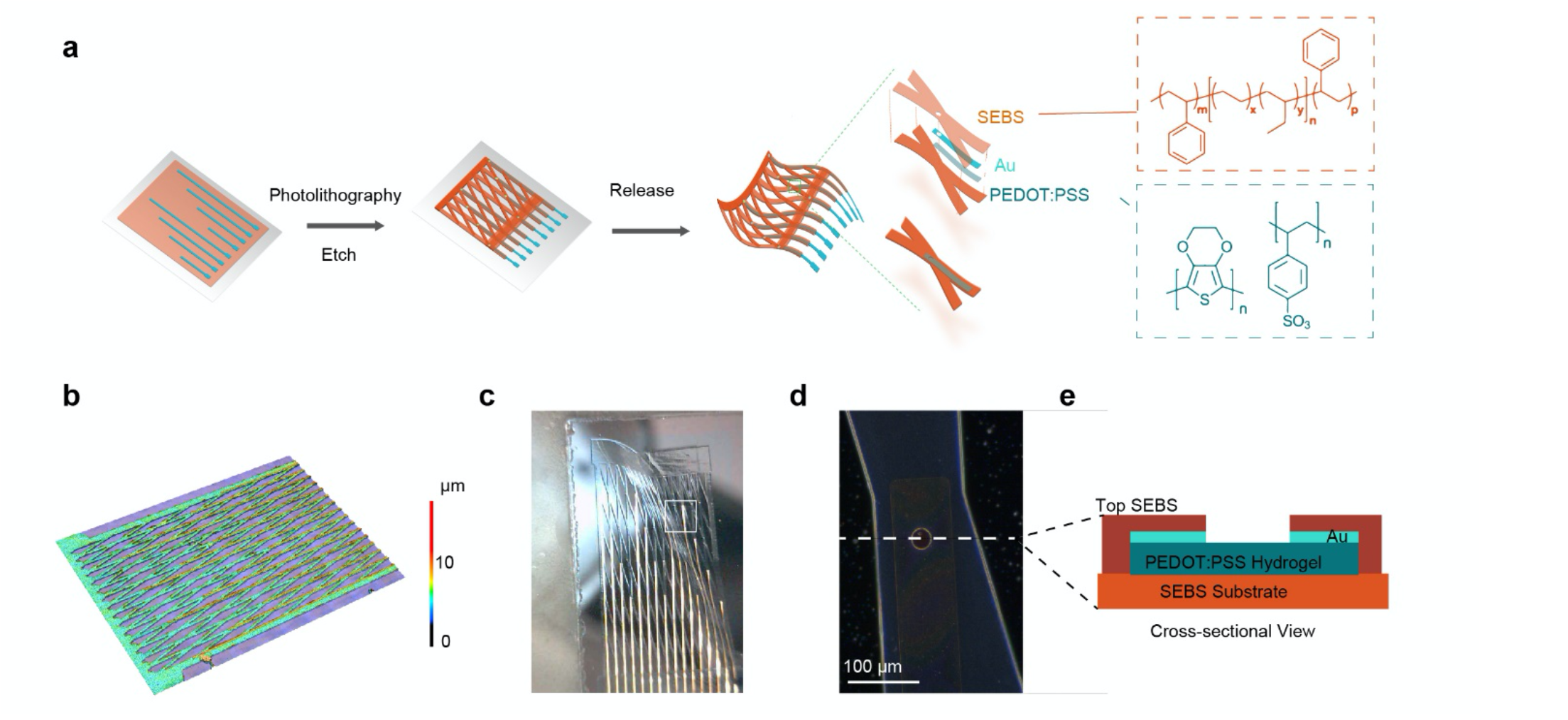
Mesh Materials, Structure, and Fabrication. (**a**) An illustration of the mesh and its fabrication process. The stretchable SEBS and PEDOT:PSS materials are patterned into a mesh shape on a silicon wafer with a sacrificial dextran layer and released using an aqueous solution. (**b**) Height mapping of the mesh indicates that it has an overall thickness of 5 μm. (**c**) A photograph of a mesh electrode after being released from its silicon wafer, demonstrating its flexibility. (**d**) A micrograph of a single electrode on the mesh.

**Figure 2:**
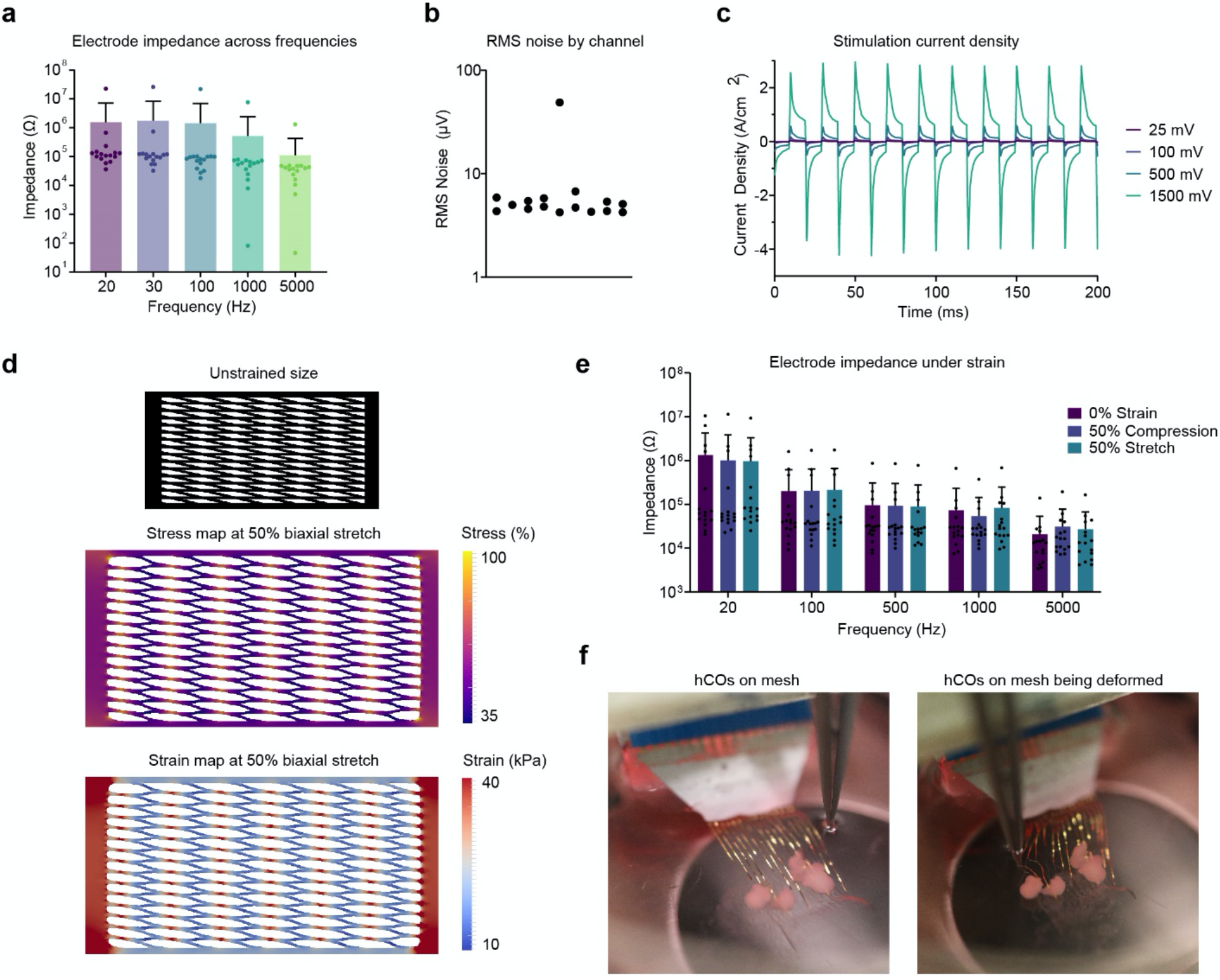
Mesh Characterization. (**a**) Measurements of the impedance of all 16 electrodes in a single mesh at varying frequencies. Error bars are ± SD. **(b)** RMS noise of all 16 electrodes in a single mesh in PBS. (**c**) The measured current density through a mesh electrode as square waves of varying voltage are applied. (**d**) Simulated stress and strain maps of the mesh under biaxial stretch. The black mesh indicates the relative size of the unstretched mesh, and the two maps indicate the local stress and strain, showing that the strain is concentrated at the junctions of the mesh.**(e)** Measurements of the impedance of all 16 electrodes under compression and stretch. Error bars are ± SD. Within each frequency, the differences between strain groups are not statistically significant.(**f**) Measurements of the impedance of all 16 electrodes in a single mesh at varying frequencies under 50% stretch and 50% compression. Error bars are ± SD. (**g**) Photographs of a mesh electrode with organoids being manipulated by metal forceps. The left image is of an undisturbed mesh, and the right image is the mesh being deformed by the forceps.

To culture hCOs on the mesh electrode, we built a hinged-lid system from a 60-mm ultra-low attachment culture plate. Using a laser cutter, we cut the lid twice: first cutting a slot in the lid for electrical contacts, and then cutting fully across the lid to separate it into two parts. We placed sterile tape across the larger cut and threaded the released mesh through the slot. This design allows electrical contacts to be made from the top of the dish, away from the media (**Fig. 3a**). Additionally, the culture media and the hCO could be accessed by opening the front of the plate, leaving the mesh itself undisturbed (**Fig. 3a**). To prepare the mesh for hCO seeding, we placed an autoclaved teflon ring on top of a geltrex-coated mesh. Three to five hCOs at day 10 of differentiation were placed inside the ring. The weight of the teflon ring minimized mechanical motion of the mesh and served to corral the floating hCOs over the mesh until they adhered. This assembly was carefully returned to the incubator, where the hCO would attach over 48 hours. After the initial attachment, the teflon ring was removed, allowing the mesh-organoid assembly to float freely in the culture medium.

**Figure 3:**
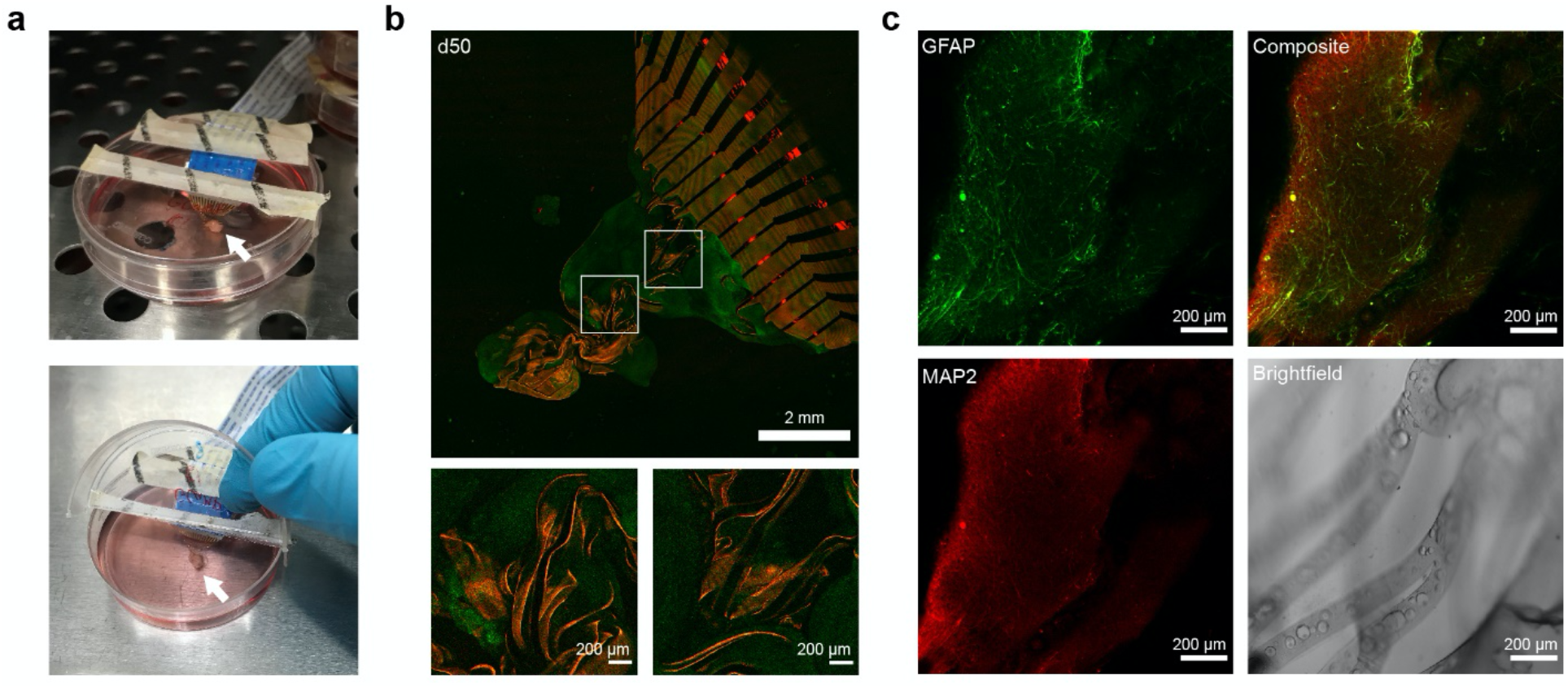
Organoid Integration. (**a**) Photographs of the hinged lid dish used to culture hCO on the mesh. The upper image shows the mesh on the hCO inside the incubator, and the lower image shows the lid opened for media changes. The hCO growing on the mesh is indicated by a white arrow, and the flat flexible cable is the white cable with printed text.(**b**) A TUBA1B-mEGFP hCO (green) growing on a rhodamine-doped mesh (red) 50 days after seeding at differentiation day 10. Both inset regions and overall image are maximum projections of a confocal z-stack. White boxes indicate the inset regions. (**c**) A CUBIC-cleared organoid grown on a dye-free mesh 114 days after seeding at differentiation day 32, stained for the glial marker GFAP (green) and the neuronal marker MAP2 (red). The composite image is a maximum projection of a confocal z-stack, and the insets are single slices from the z-stack.

We visualized the integration of hCOs into the mesh using rhodamine doped into the top layer of the SEBS. We seeded these fluorescent meshes with hCOs derived from TUBA1B-mEGFP hiPSCs, which have been genetically engineered to express GFP-tagged α-tubulin (Coriell). Over time, the hCO grew into and integrated with the mesh. By day 50 after plating, the organoid had grown and become well integrated into the mesh material, as shown by eGFP-positive organoids encapsulating the red mesh lines (**Fig. 3b**).

In addition to live imaging, we performed intact hCO staining and tissue clearing. For this purpose, we used meshes without rhodamine labeling. More specifically, we seeded an hCO on a mesh at day 32 of differentiation, and fixed while integrated with the mesh at day 146 of differentiation, for a total of 114 days on the mesh. After fixation, the hCO was stained for MAP2– a marker of neuronal dendrites, and GFAP– a marker for glial cells. We then cleared the hCO according to the CUBIC protocol^32^. This tissue clearing approach showed that the hCO is well-integrated into the mesh (**Fig. 2c**). Upon examination of individual z-slices, we found no evidence of increased GFAP signal near the mesh material, suggesting that these meshes inside organoids did not necessarily elicit obvious glial responses. Likewise, there was little observable difference in the MAP2 channel in areas that are close to the mesh and areas that are far away. In the CUBIC-cleared mesh, we observed some green fluorescence artifacts that correspond to mesh traces, but these do not appear to be signal from cells. Taken together, our results suggest that the mesh material and design is compatible with hCO integration and growth.

Having established that the mesh is biocompatible and integrates with hCOs, we then tested the electrical interface between the organoid and the mesh electrode using electrical stimulation. First, we determined the minimum stimulation parameters necessary to stimulate the hCO. For these initial experiments, we inserted a bipolar stereotrode into a day 129 hCO that expressed the genetically encoded calcium indicator GCaMP6f. We then performed simultaneous electrical stimulation and confocal imaging (**Fig. S3a**). We screened different stimulation parameters, including the current amplitude, the pulse width, the number of pulses, and the pulse train frequency. To understand the localization of stimulation, we analyzed two regions of interest (ROIs): one drawn around the region immediately surrounding the stereotrode location (ROI 1), and a region several hundred microns away (ROI 2). First, we tested stimulation at varying currents. We fired a pulse train of 100 pulses at 250 Hz at amplitudes ranging from 50 μA to 5 μA, and found that 50 μA stimulations evoked calcium signals at both ROIs, 20 μA and 10 μA only evoked a signal in the proximal ROI, and 5 μA failed to elicit any response (**Fig. S3b**). Next, we tested the effect of pulse train frequency. Using a stimulation amplitude of 20 μA and using 100 pulses, we tested stimulation at 250 Hz and 50 Hz. We found that stimulating at 50 Hz was able to elicit a response in the distal ROI, whereas 250 Hz did not (**Fig. S3c**). Finally, using 20 μA and 50 Hz, we tested the effects of changing the pulse number on stimulation, varying the number of pulses per train from 100 pulses to 1 pulse, finding that as few as 5 pulses per pulse train was sufficient to elicit a response in the near ROI (**Fig. S3d**).

Using this information, we determined whether the meshes were capable of stimulating hCOs. After an organoid was placed on the mesh at 10 days of differentiation and cultured on the mesh for 80 days (90 days of differentiation), we infected the integrated organoid with an AAV encoding a calcium indicator GCaMP6f. We performed the same stimulation experiment through the mesh electrodes using the optimized parameters obtained from the sterotrodes. To do so, we connected the mesh organoid to the Intan RHS controller (**Fig. 4a**). To validate the mesh’s integration with the organoid, we measured the open circuit potential of each electrode using a potentiostat. When compared to a calibration curve measured in standard pH solutions, the open circuit potential can be used to estimate the local pH. According to these measurements, the pH at different electrode locations varied between 6.4–7.2 with a mean value of 6.7 ± 0.4 (**Fig. 4b**). These slightly acidic values are additional evidence that the electrodes are indeed embedded inside the organoid, where the local environment may be more acidic due to metabolic byproducts.

**Figure 4:**
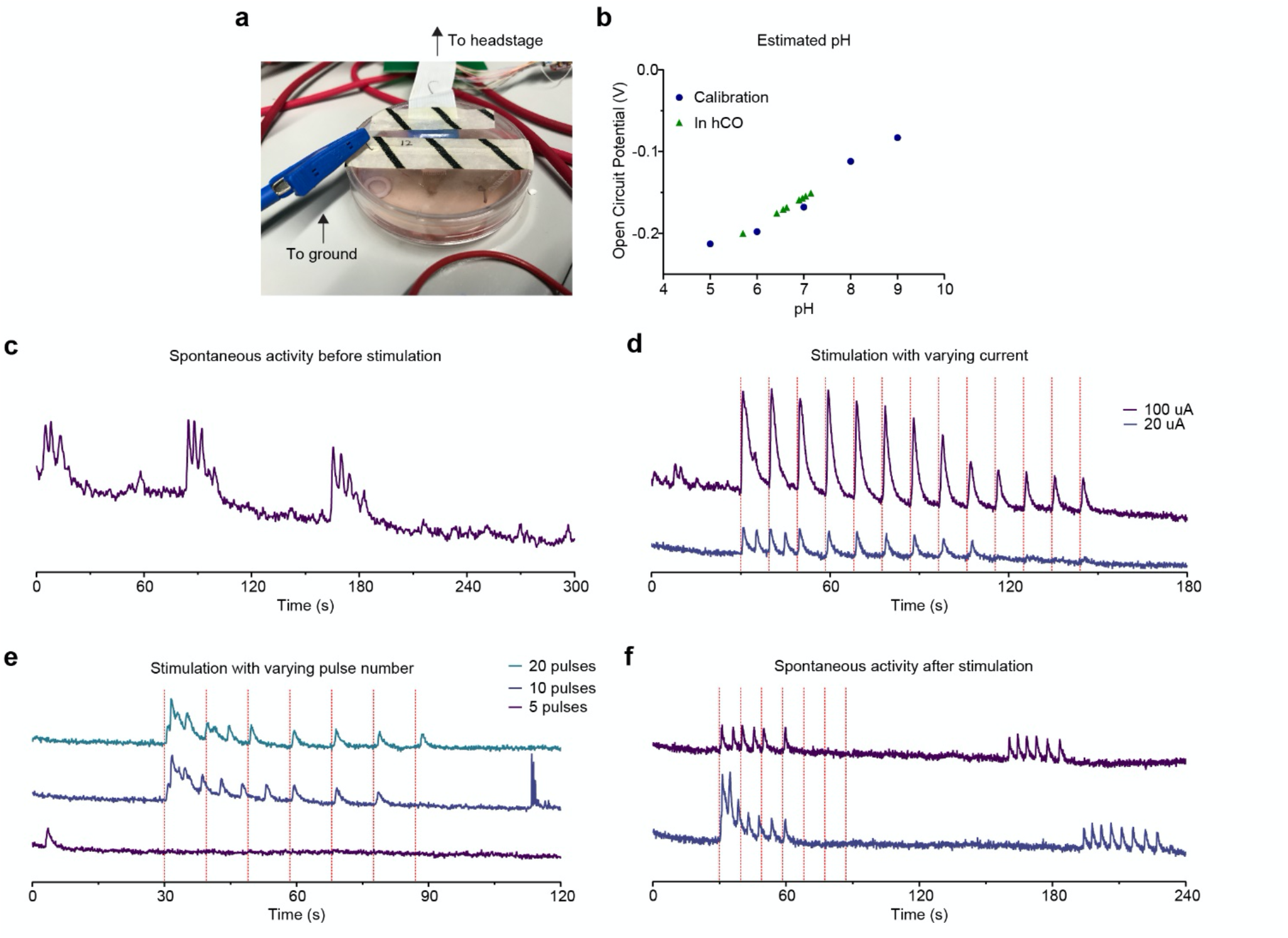
Electrical Interfacing Between the Mesh and Organoid. (**a**) A photograph of how the mesh electrode is connected to the Stim/Recording controller. The white FFC is connected to the adapter and headstage, and a platinum wire is inserted through the tape at the hinge. (**b**) pH estimates based on the measured open circuit potential at each electrode in an organoid. pH levels were calibrated by adjusting the pH of a reference PBS solution. **(C)** A GCaMP7f intensity plot showing spontaneous calcium activity in a d90 hCO before stimulation. Intensity was calculated by averaging the intensity of the entire frame at each timepoint. (**d**) A GCaMP intensity plot showing calcium spikes in response to stimulation of varying currents applied to the mesh. Each vertical red line indicates one pulse train consisting of 100 pulses at 100 Hz at the indicated current. (**e**) A GCaMP intensity plot showing calcium spikes in response to stimulation applied to the mesh. Each vertical red line indicates one pulse train consisting of pulses with an amplitude of 20 μA and a frequency of 100 Hz, with the indicated number of pulses. (**f**) A GCaMP intensity plot showing extended imaging after stimulation. Each vertical red line indicates one pulse train consisting of 10 pulses with an amplitude of 20 μA and a frequency of 100 Hz.

We then measured the calcium responses of the hCO. First, we measured a baseline of activity in the hCO, and observed that there were several bursts of spontaneous activity (**Fig. 4c**). We then tested the observation that different stimulation amplitudes result in different degrees of calcium signals. We triggered a stimulation through all 16 channels of the mesh electrode at 100 pulses and 100 Hz at 100 μA and 20 μA, measuring the average intensity of the entire frame. The 100 μA stimulations elicited a larger response than the 20 μA stimulations, and were able to continue eliciting responses as more pulse trains were fired (**Fig. 4d**). The differences in the responses can be partially explained by differences in the localization of the stimulation. Dividing the timelapse into 4 different spatial regions (**Fig. S4a**) showed that the 100 μA stimulation drives a calcium signal in three of the four regions (**Fig. S4b**), whereas the 20 μA stimulation affects only one area (**Fig. S4c**). We then tested the effect of varying the number of pulses per train on stimulation. Consistent with our observations using the bipolar stereotrode, we were able to elicit calcium activity using 20 or 10 pulses per train at 20 μA, but 5 pulses were insufficient (**Fig. 4f**). Finally, after the stimulation experiments were performed, we imaged for several minutes after stimulating with 10 pulses at 100 Hz and 20 μA, and observed spontaneous bursts of activity several minutes after stimulation. We noted that these post-stimulation spontaneous spikes were well-separated, with each spike returning to the baseline before the next spike, in contrast to the spontaneous spikes observed before stimulation, which were more tightly clustered.

## Discussion

Here, we have developed a novel stretchable soft mesh electrode system. While previous mesh electrode systems use non-deformable flexible materials, the SEBS/PEDOT electrodes are intrinsically stretchable and can sustain substantial strain without affecting its electrical performance. These meshes can establish a stable, 3D long-term interface with hCOs for more than 3 months, as the organoid-mesh hybrid grows to several millimeters in size. As a result of this integration, we were able to apply electrical stimulations throughout the hCO, as demonstrated by calcium imaging during stimulation. The necessary stimulation parameters and results were consistent with those using a more traditional bipolar stereotrode.

Our system demonstrates the potential of a long-term electrophysiological interfaces with hCOs. Because the mesh system is well integrated into the 3D structure of the organoid, experiments and measurements can be performed on the same hCO at multiple timepoints, as the stretchability of its materials enables the mesh electrode to adapt as the 3D culture grows. This stretchable mesh system could apply repeated chronic stimulations to organoids over time. Additional work is needed to improve the scalability of the system and enable reliable recording. Simultaneous recording and stimulation would enable the active modulation of electrical activity over extended periods and in models of disease. In addition to direct electrical measurements, factors such as pH and neurotransmitter concentrations could be monitored over time. Thus, this mesh electrode system potentially opens new avenues for studying self-organizing human cellular systems.

## Author Contributions

All authors contributed to the study conception and design. TL performed the cell culture and imaging and drafted the manuscript. YL designed, fabricated, and characterized the mesh electrodes. TL and YL designed the culture setup, performed the stimulation experiments, and performed data analysis. CF and LB performed the stretching simulations. ZB, BC, and SP oversaw the project.

## Additional Information

Supplementary information is available for this paper. Correspondence and requests for materials should be addressed to Sergiu P. PaŞca, Bianxiao Cui or Zhenan Bao.

## Materials and Methods

### Mesh Fabrication

PEDOT:PSS (PH1000) and the ionic liquid (IL, 4-(3-Butyl-1-imidazolio)-1-butanesulfonic acid triflate) were purchased from Clevios and Santa Cruz Biotechnology, respectively. About 50 wt% of IL vs. PEDOT:PSS was added to PEDOT:PSS aqueous solution (0.172 g in 15 ml of PH1000 solution) and stirred vigorously for 20 min. The PEDOT:PSS/IL aqueous mixture was then filtered by a 0.45 µm syringe filter. A dextran (Sigma-Aldrich #09184) aqueous solution (5wt%) was spin coated on a Si wafer as the sacrificial layer. SEBS (20% in toluene) was spin coated at a speed of 4000 rpm for 1 min, and followed by annealing at 100 °C for 10 min. The substrate was then oxygen plasma (Technics Micro-RIE Series 800) treated at 150 W for 1 min. The PEDOT:PSS/IL aqueous mixture was drop cast or spin coated (2000 rpm, 1 min) on the oxygen plasma treated PFPE-DMA layer and baked at 110 °C for 20 min. 40 nm of Au was subsequently deposited by e-beam evaporation (KJ Lesker evaporator) or thermal evaporator. S1805 photoresist was spin coated on Au at 4000 rpm for 45 s and exposed with mask aligner (Quintel Q4000) for 3 s at a power of 10 mW/cm2 after 115 °C pre-expose baking (1 min). The exposed photoresist was developed in the MF-319 developer (Micropost, Shipley) for 1 min. Micropatterns of S1805 were transferred to Au by etching using Argon ion milling (Intlvac Nanoquest Ion Mill, bias voltage 100 V) for 1 min. Inductively coupled plasma etch system (PlasmaTherm Oxide Etcher600 W, 80 sccm) or oxygen plasma (March PX-250 Plasma Asher, 300 W, 8 sccm) was used to etch PEDOT:PSS/IL ion gel and remove the photoresist. SEBS (20% in acetone) was spin coated at a speed of 1000 rpm for 1 min, and followed by annealing at 100 °C for 10 min. 60 nm of Al was subsequently deposited by thermal evaporator. S1805 photoresist was spin coated on Al at 4000 rpm for 45 s and exposed with mask aligner (Quintel Q4000) for 3 s at a power of 10 mW/cm2 after 115 °C pre-expose baking (1 min). The exposed photoresist was developed in the MF-319 developer (Micropost, Shipley) for 1 min. Micropatterns of S1805 were transferred to Al by etching using Argon ion milling (Intlvac Nanoquest Ion Mill, bias voltage 100 V) for 1 min. Inductively coupled plasma etch system (PlasmaTherm Oxide Etcher600 W, 80 sccm) was used to etch the exposed electrode and the SEBS. Then, the Au on top of PEDOT:PSS in exposed microelectrode layer was removed by argon ion milling etching. The top Al mask was etched by FeCl3 for 5 min. Anisotropic conductive tape (3M 9703) was applied directly onto the gold pad with 1 mm pitch, at ambient temperature. The FFC jumper cables were connected to the top side of the anisotropic conductive tape via gentle press. Lastly, the free-standing device was obtained by soaking in water for at least 1 day and subsequently the entire film was released from the dextran sacrificial layer. The substrate was then oxygen plasma (Technics Micro-RIE Series 800) treated at 150 W for 15s prior soaking in geltrex and incubated in 37C overnight.

### Electrical Characterization of the Mesh

An Intan RHS Stim/Recording System was used to interface with the mesh electrodes. The FFC of the mesh assembly was connected to a breakout board containing an FFC connector (NHD-FFC16-1, Newhaven Display) leading to 16 through-holes. A 16-pin wire adapter (Intan B7600) was soldered to these through-holes, and connected to a 16-channel Stim/Recording Headstage (Intan M4016). Impedance, noise, and stimulation measurements were performed through the Intan RHS Data Acquisition Software. For impedance characterization during stretch, a tweezer with a fine tip was used to pull the mesh, which is floating in culture media, and the amplitude of the impedance were measured at different strains.

### Mesh Mechanical Simulations

The material styrene-ethylene-butylene-styrene (SEBS) used to make the mesh-electrode-array has been shown to behave in the hyperelastic regime (https://doi.org/10.1007/s10853-017-0991-z) The simulation takes the CAD-design of the mesh-electrode-array’s envelope and neglects the presence of the gold layer, which is only 40 nanometers thin. To simulate the mechanical deformation, we applied the equations of hyperelasticity in the Mooney-Rivlin formulation of the mechanical energy functional (https://doi.org/10.1098/rsta.1948.0024). A Python package for solving finite-element-analysis problems, FEniCS (https://doi.org/10.11588/ans.2015.100.20553), was used to compute the deformation of the mesh. The package contains a native meshing algorithm to discretize the CAD-design into a triangular mesh to evaluate the necessary differential equations. For boundary conditions, we use Dirichlet boundary conditions and impose a 50 percent elongation of the mesh in the x and y plane. The code to generate the discretization and compute the deformation is freely available and can be reproduced at github.com/bcuilab/stretching_simulation].

### Generation of hCO

Human cortical organoids (hCO), also known as cortical spheroids, were derived according to our previously established methods^15^. Briefly, hiPSC cultured on DR4 feeders were cultured using the following culture medium: DMEM/F12 (1:1) (Invitrogen) containing 20% KnockOut Serum (Invitrogen), 1 mM non-essential amino acids (Invitrogen, 1:100), GlutaMax (Invitrogen, 1:100), 0.1 nM β-mercaptoethanol (Sigma-Aldrich), 100 U/ml penicillin and 100 μg/ml streptomycin (Invitrogen) and 10 ng/ml FGF2 (R&D Systems) Human pluripotent stem cell colonies were lifted off using dispase (Invitrogen: 17105-041; 0.7 mg/ml) and transferred to ultra-low-attachment 100 mm plastic plates (Corning). For the first 24 h, the colonies were maintained in stem cell medium supplemented with Y-27632 (EMD Chemicals). Dorsomorphin (Sigma, 5 μM) and SB-431542 (Tocris, 10 μM) until day 5. On day 6, the media was changed to the following medium: Neurobasal-A (Gibco 10888022), B-27 without vitamin A (Gibco 12587010), GlutaMax (Gibco 35050061, 1:100), and 100 U/ml penicillin and 100 μl streptomycin (Gibco). The media was supplemented with 20 ng/ml FGF2 (R&D Systems) and 20 ng/ml EGF (R&D Systems) for 19 days with daily medium change in the first 10 days, and every other day for the subsequent 9 days. At day 25, FGF2 and EGF were replaced with 20 ng/ml BDNF (Peprotech) and 20 ng/ml NT3 (Peprotech) At day 43 onwards, growth factors were removed and medium was changed every 4 days.

### hCO integration

Hinged petri dishes were prepared using 60 mm ultra-low attachment plates (Corning). Batches of the 10 mm-plate lids were cut using a laser cutter (Epilog Fusion M2). Two lines were cut, dividing the lid into thirds: one line did not reach to the edges of the lid, creating a slot, and the other line was cut fully across the lid, splitting it in two.

To prepare meshes for hCO integration, mesh-FFC assemblies were first sterilized, by dipping in 70% ethanol and drying in the tissue culture hood. Sterilized meshes were placed mesh-side down in a 50 mL falcon tube. A coating solution was prepared with a 1:100 dilution of Geltrex (Thermo Fisher A1413201) diluted in PBS. This coating solution was added to the falcon tube such that only the mesh and wafer were submerged in liquid, and placed in the incubator for 24 h. This process both coated the mesh and dissolved the dextran sacrificial layer, releasing the mesh.

After incubation, the meshes were transferred to a 100 mm cell culture plate filled with PBS. If the meshes were not released from the wafer at this point, additional gentle force was applied to separate the wafer.

Meanwhile, the cut lids were sterilized with ethanol, and a length of autoclave tape was placed in a pipette tip box and autoclaved for sterilization. The two cut pieces of each lid were re-aligned and connected with a strip of autoclave tape.

The bottoms of the petri dishes were filled with cell culture media. The FFC, with the mesh on the end, was threaded through the laser-cut slot in the lid, and the lid and mesh assembly were placed on the bottom of the petri dish. The height of the mesh in the dish was adjusted manually such that the mesh floated freely in the solution, but the FFC contact pads were not submerged. After setting this height, a strip of autoclave tape was placed over the slot. This served to seal any gaps in the lid, and to fix the position of the mesh.

Organoids at day 10 of differentiation were then seeded on the mesh. If the mesh was not freely floating, it would be spread out using a pipette tip. A pre-autoclaved teflon ring (55D Durometer, Round, White, 1/2” ID, 5/8” OD, 1/16” Width, Sterling Seal & Supply, Inc. (STCC)) was placed on top of the mesh using forceps. This ring served to corral floating organoids, and to hold the mesh down during seeding. 3-5 organoids at d10 in differentiation were placed on top of the mesh inside the ring, and the dish was carefully transferred to the cell culture incubator. Organoid attachment was observed 24 h after seeding. If the organoids did not attach to the mesh or had floated elsewhere in the mesh, they would be re-placed within the ring until they successfully adhered to the mesh. After the organoids were successfully adhered, the teflon ring was discarded.

Media changes were performed according to the original protocol. To access the media, only the front half of the lid was flipped upward, using the autoclave tape as a hinge. This allows media changes and access to the organoid without disturbing the FFC or the mesh.

### hCO Imaging and Clearing

Tissue clearing was performed according to the CUBIC protocol^32^. Briefly, hCO were fixed with 4% paraformaldehyde at 4 C overnight, incubated with CUBIC-L Tissue Clearing Reagent (TCI T3740) for 48 hours at 37 C, with the CUBIC-L refreshed at 24 hours. Organoids were washed with PBS overnight, then incubated in MAP2 and GFAP primary antibodies dissolved in blocking solution (0.2% Triton-X and 3% Normal Goat Serum in PBS) for 48 hours at 37 C. They were then washed with PBS, with 3 washes of 2 hr and 1 overnight incubation at room temperature. Organoids were then incubated with secondary antibodies (Alexa Guinea Pig 647 and Alexa Rabbit 568, 1:500 dilutions) in blocking solution for 48 hours at 37 C. After secondary antibody staining, organoids were washed with PBS, with 3 washes of 2 hr and 1 overnight incubation at room temperature, and exposed to a pre-treatment of 1:1 water and CUBIC-R+ (TCI T3741) for 6 hours at room temperature. After pre-treatment, stained organoids were incubated in 100% CUBIC-R+ for 48 hours for clearing. Cleared samples were imaged using a Leica SP8 laser-scanning confocal microscope.

### pH Measurements

pH measurement was calculated by using open circuit potential. Potentiostat (Palmsens) was used to read out the open circuit potential between the mesh electrode and reference electrode. The mesh electrodes were first calibrated in standard pH solution at various pH. After successful integration between the organoid and the mesh electronics, the open circuit potential was measured to quantify the pH environment within the organoid. The calibration curve was used to estimate the pH environment within the organoid.

### Electrical Stimulation

Electrical stimulation was performed through the Intan RHS 16-channel headstage and Intan Stim/Recording system.

Organoids were infected with a GCaMP7f-AAV (pAAV.CAG.GCaMP6f.WPRE.SV40, Addgene) which was delivered at day 119. The virus was added to the mesh culture dish, and the media was exchanged 24h after adding the virus. The organoids were then maintained normally for the defined interval (10 or 14 days) before imaging.

For probe stimulation, a bipolar platinum/iridium stimulation probe was used (Microprobes PI2ST30.1A10). One of the probe plugs was connected to the RHS headstage by an alligator clip, and the other end of the wire was clipped to a stripped wire adapter (Intan B7600). The other probe plug was connected via alligator clips and a jumper cable to the GND port of the headstage. The stimulation probe was taped onto a micromanipulator (Sutter MP-225) and placed on the stage of Leica SP8 laser-scanning confocal microscope with a stage-top incubator for CO2 and temperature control (OKO Lab)

For mesh stimulation, GCaMP7f-expressing organoids cultured on meshes were placed on the microscope stage and connected to the 16-channel RHS headstage as described above.

In both cases, stimulation parameters were set in the Intan RHS software, and imaging and stimulation timepoints were set using the Leica LASX software in Live Data Mode. The Trigger Unit of the confocal microscope was connected to the digital input of the Intan Sim/Recording system, so that stimulations would be triggered on particular frames of the timelapse. In addition to the parameters discussed in the results, each stimulation was a biphasic stimulation, cathodic first, with a 200μs phase duration. The intensity plots for calcium imaging were computed by calculating the average intensity of the entire frame for each timestamp, or by calculating the average intensity of the indicated region of interest.

**Figure S1:**
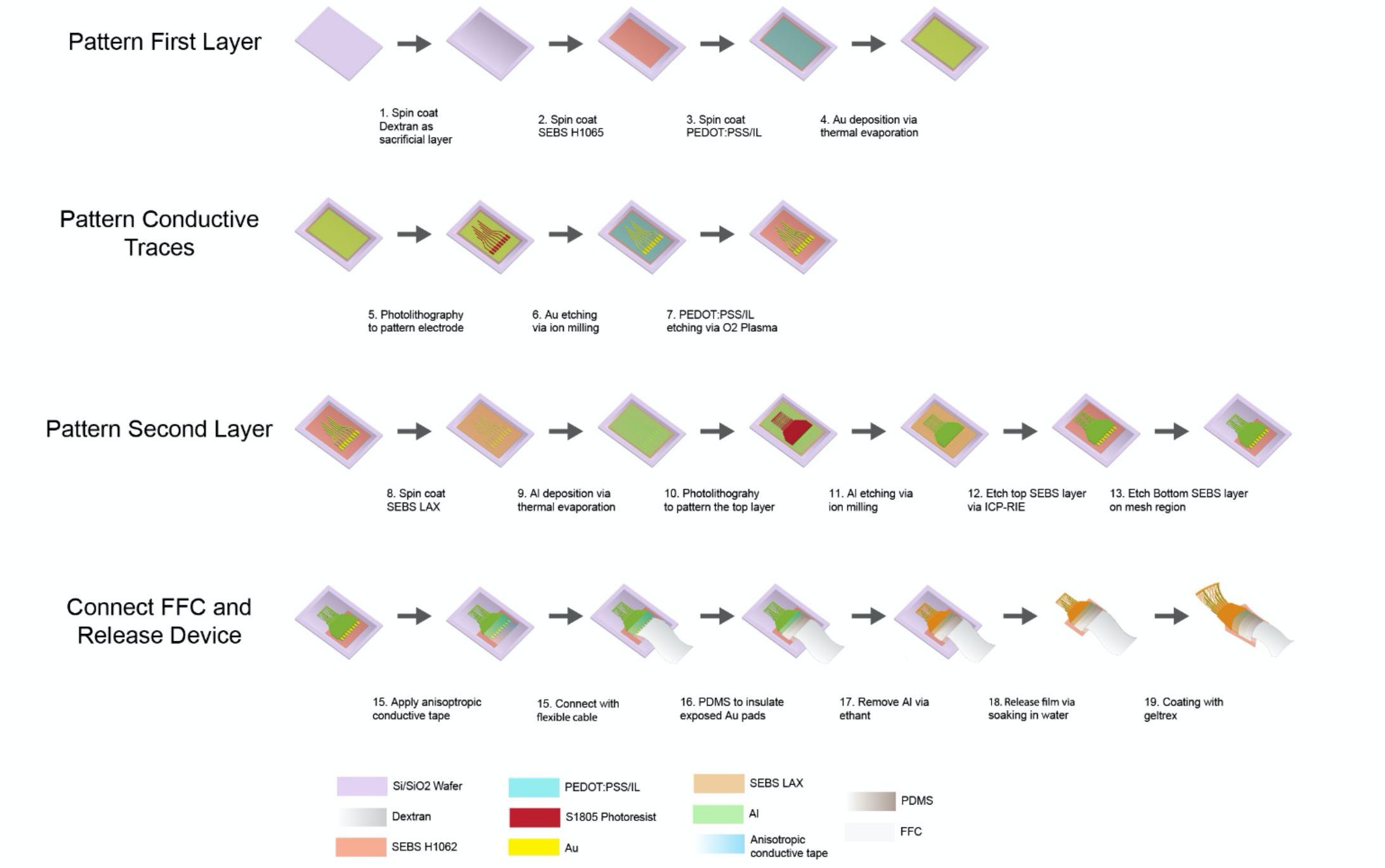
Mesh Fabrication. A schematic representation of the mesh fabrication process. First, we spin-coated a sacrificial layer of dextran onto a silicon wafer, followed by SEBS, PEDOT:PSS ionic liquid, and gold. We then patterned these layers by sequential photolithography and etching. We patterned and etched a second layer of SEBS to complete the insulation and performed additional etching to expose the electrodes and connection pads. After the mesh was fully patterned, we connected the exposed pads to an FFC using anisotropic conductive tape, encapsulated the connection with PDMS, and soaked the entire assembly in water to release it from the silicon wafer.

**Figure S2:**
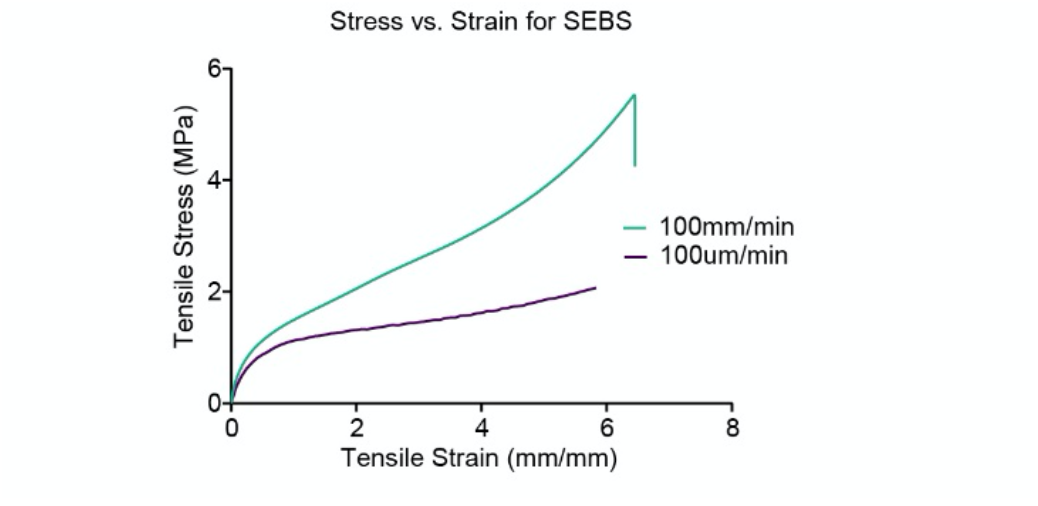
SEBS Sheet Stress-Strain Curve. The measured stress-strain curve of a thin film of SEBS, without a mesh structure, under two different stretch rates. The lower stress at lower stretch rates demonstrates the viscoelasticity of SEBS.

**Figure S3:**
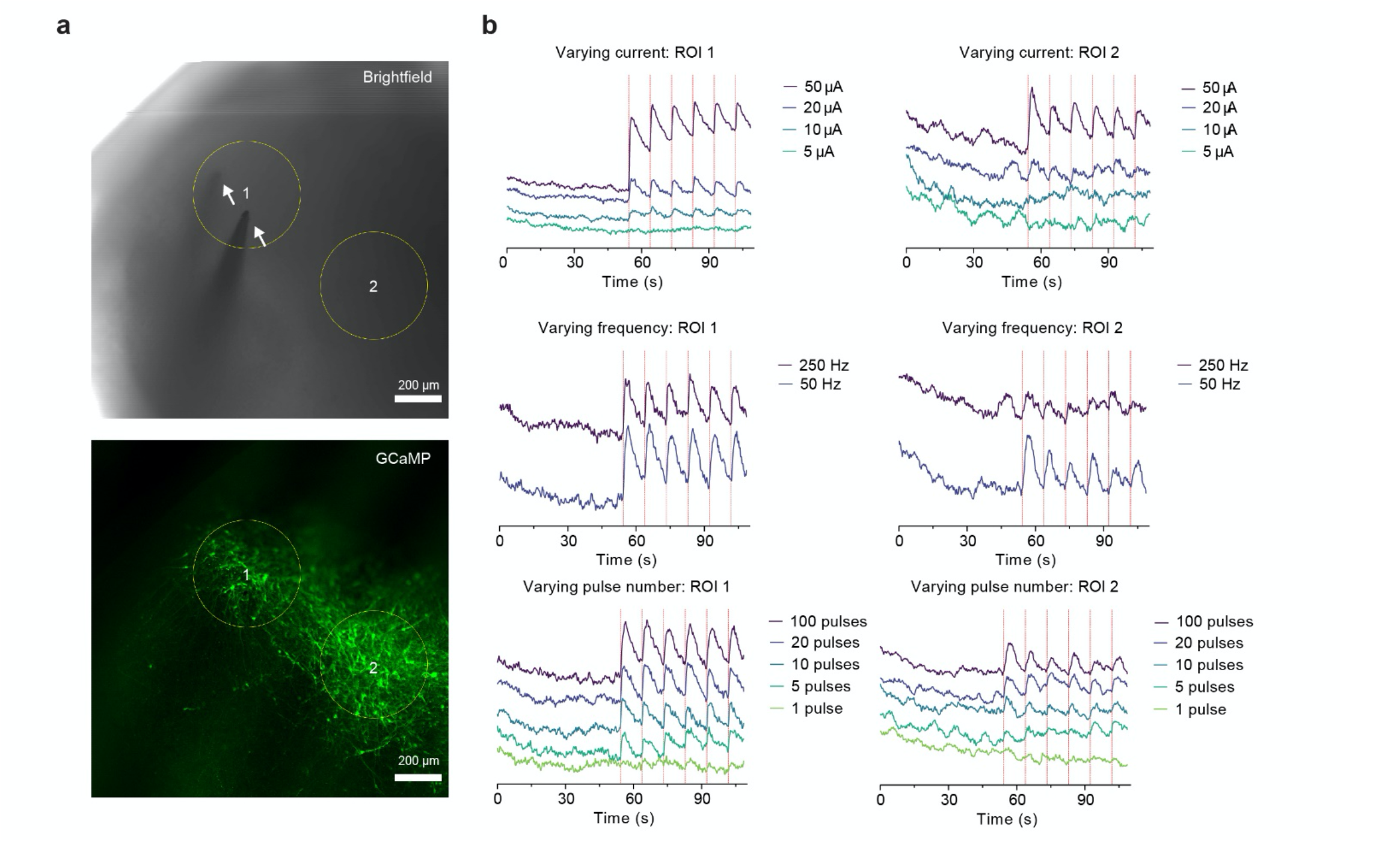
Probe Stimulation. (**a**) Microscope images of stereotrode stimulation. Images were generated by the average projection of one time-lapse. White arrows in the brightfield image indicate the locations of the two points of the stereotrode. Yellow circles indicate the two ROIs under analysis. (**b**) GCaMP7f intensity plots of the two ROIs of the timelapse under different stimulation conditions.

**Figure S4:**
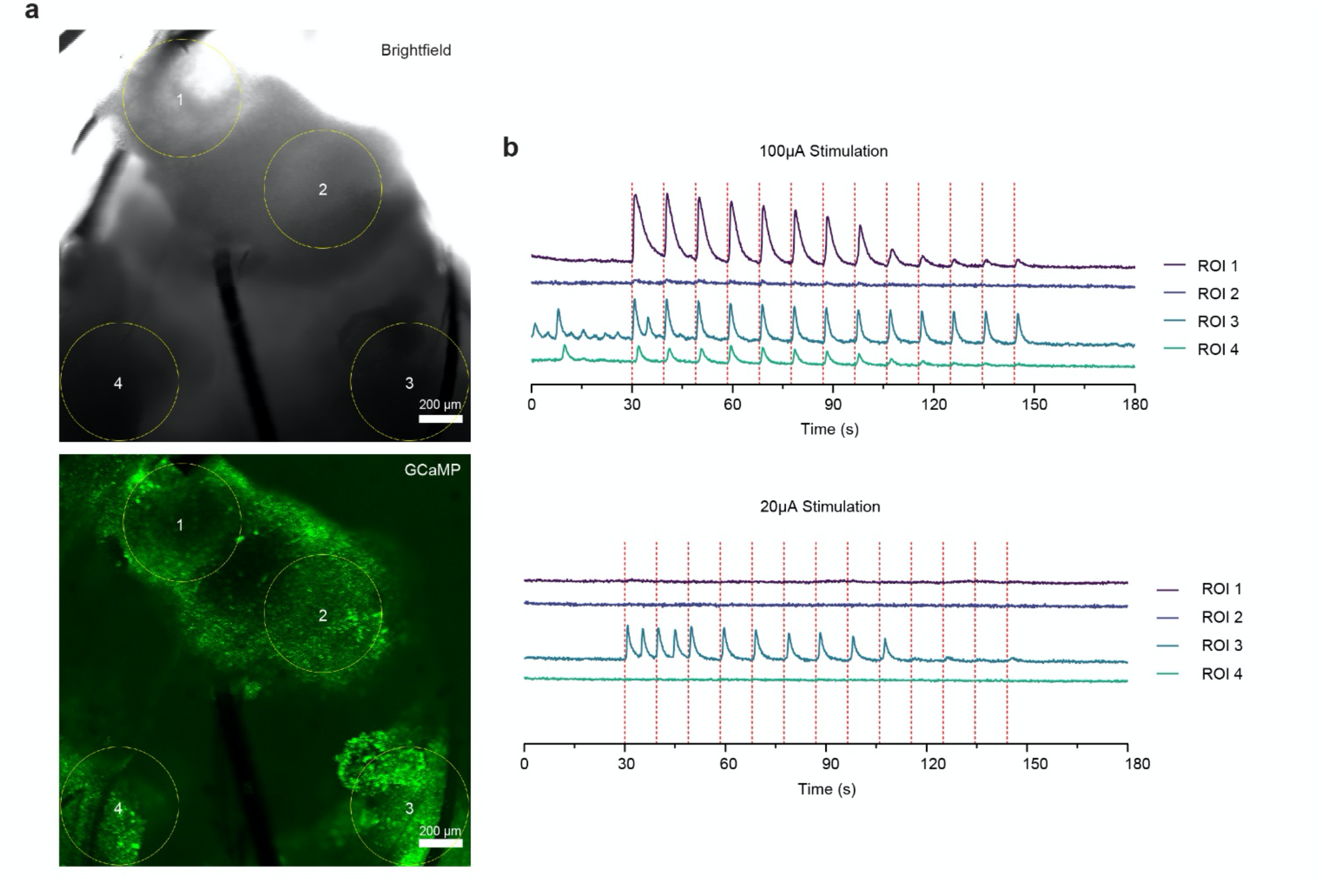
Localization of Stimulation. (**a**) Microscope images of mesh stimulation. Images were generated by the average projection of one time lapse. Yellow circles indicate the four ROIs under analysis. (**b**) GCaMP intensity plots of the four ROIs of the timelapse in response to stimulation at 100 μA and 20 μA.

